# A single amino acid substitution in PB1 of pandemic H1N1 with A/Ann Arbor/6/60 master donor virus mutations as a novel live-attenuated influenza virus vaccine

**DOI:** 10.1101/2022.01.07.475442

**Authors:** Aitor Nogales, John Steel, Wen-Chun Liu, Anice C. Lowen, Laura Rodriguez, Kevin Chiem, Andrew Cox, Adolfo García-Sastre, Randy A. Albrecht, Stephen Dewhurst, Luis Martínez-Sobrido

## Abstract

Influenza A viruses (IAV) remain emerging threats to human public health. Live-attenuated influenza vaccines (LAIV) are one of the most effective prophylactic options to prevent disease caused by influenza infections. However, licensed LAIV remain restricted for use in 2- to 49-year old healthy and non-pregnant people. Therefore, development of LAIV with increased safety, immunogenicity, and protective efficacy is highly desired. The United States (U.S.) licensed LAIV is based on the master donor virus (MDV) A/Ann Arbor/6/60 H2N2 backbone, which was generated by adaptation of the virus to growth at low temperatures. Introducing the genetic signature of the U.S. MDV into the backbone of other IAV strains resulted in varying levels of attenuation. While the U.S. MDV mutations conferred an attenuated phenotype to other IAV strains, the same amino acid changes did not significantly attenuate the pandemic A/California/04/09 H1N1 (pH1N1) strain. To attenuate pH1N1, we replaced the conserved leucine at position 319 with glutamine (L319Q) in PB1 and analyzed the *in vitro* and *in vivo* properties of pH1N1 viruses containing either PB1 _L319Q_ alone or in combination with the U.S. MDV mutations using two animal models of influenza infection and transmission, ferrets and guinea pigs. Our results demonstrated that L319Q substitution in the pH1N1 PB1 alone or in combination with the mutations of the U.S. MDV resulted in reduced pathogenicity (ferrets) and transmission (guinea pigs), and an enhanced temperature sensitive phenotype. These results demonstrate the feasibility of generating an attenuated MDV based on the backbone of a contemporary pH1N1 IAV strain.

**IMPORTANCE:** Vaccination represents the most effective strategy to reduce the impact of seasonal IAV infections. Although LAIV are superior in inducing protection and sterilizing immunity, they are not recommended for many individuals who are at high risk for severe disease. Thus, development of safer and more effective LAIV are needed. A concern with the current MDV used to generate the U.S. licensed LAIV is that it is based on a virus isolated in 1960. Moreover, mutations that confer the temperature sensitive, cold-adapted, and attenuated phenotype of the U.S. MDV resulted in low level of attenuation in the contemporary pandemic A/California/04/09 H1N1 (pH1N1). Here, we show that introduction of PB1 _L319Q_ substitution, alone or in combination with the U.S. MDV mutations, resulted in pH1N1 attenuation. These findings support the development of a novel LAIV MDV based on a contemporary pH1N1 strain as a medical countermeasure against currently circulating H1N1 IAV.

## INTRODUCTION

Influenza A viruses (IAV) are enveloped viruses that contain a genome comprising eight single-stranded negative-sense RNA segments (1, 2). IAV are classified based on the antigenic properties of the two viral surface glycoproteins, hemagglutinin (HA) and neuraminidase (NA). Currently, only two IAV subtypes (H1N1 and H3N2) circulate in humans and result in seasonal influenza epidemics of mild to severe respiratory illness with instances of fatal outcomes (3-7). Despite global vaccination campaigns, which represent the most cost-effective strategy to prevent IAV infections (4, 5, 7-13), it is estimated that seasonal influenza infections are still responsible for approximately 4 million cases of severe disease and 500,000 deaths worldwide yearly (14-16) (https://www.who.int/news-room/fact-sheets/detail/influenza-(seasonal)). In addition, zoonotic IAV can cause sporadic pandemics of severe consequences (17-24).

There are two major types of licensed IAV vaccines for clinical application: inactivated influenza vaccines (IIV) and live-attenuated influenza vaccines (LAIV). IIV mainly induce humoral but not cell-mediated immune responses (25-34). Therefore, IIV offers limited protection if the seasonal vaccines do not antigenically match the predicted circulating IAV strains (9, 10, 13, 15). Furthermore, the immunogenicity and protective efficacy of IIV is reduced in immunocompromised patients and elderly individuals (35-37). On the other hand, LAIV have the potential to elicit broader protection because they induce both humoral and cell-mediated immune responses (5, 9, 10, 27, 38). Moreover, since LAIV are delivered nasally, they induce strong mucosal immunity at the site of infection (8, 35, 39). Finally, due to their induction of a cross-reactive cell-mediated immunity, which targets conserved viral proteins, LAIV confer more efficient protection against heterologous IAV strains, which is important in the case of antigenic mismatch between vaccine and circulating strains (9, 10, 27-34, 40-43). Unfortunately, because safety concerns, the United States (U.S.) licensed LAIV remains restricted to 2- to 49-year old healthy, non-pregnant persons. In addition, the increased levels of IAV pre-existing immunity in the >50-year old adults can also limits the LAIV uptake (4, 8, 44). As a result, many individuals who are at high risk for severe influenza are unable to receive LAIV or vaccine efficacy is poor. Therefore, the development of a novel LAIV with improved immunogenicity, safety, and protective efficacy is highly desirable.

Current LAIV leverage the temperature gradient between the human upper and lower respiratory tract (URT and LRT, respectively), and are characterized by cold-adapted (*ca*), temperature-sensitive (*ts*) and attenuated (*att*) phenotypes. LAIV replicate in the cooler URT, where they induce protective immune responses, but LAIV do not replicate in the LRT due to the elevated temperatures, which limits the potential risk of pulmonary damage and disease (5, 38, 45-47). The A/Ann Arbor/6/60 H2N2 master donor virus (MDV) used to manufacture currently licensed U.S. LAIV was generated by passaging the virus at gradually reduced temperatures, thereby selecting mutants that replicate efficiently at low temperatures (*ca*) but not at elevated temperatures (*ts*) (5, 7, 9, 10, 38, 48-50). Five mutations within the viral replicative machinery have been shown to be responsible for the *ts*, *ca*, and *att* phenotype of the U.S. A/Ann Arbor/6/60 H2N2 LAIV MDV: the viral polymerase subunits PB2 (N265S) and PB1 (K391E, D581G, and A661T); and the viral nucleoprotein, NP (D34G) (5, 38, 45, 48-50).

A concern with this U.S. MDV is that it is based on a virus isolated in 1960 and a more contemporary MDV might be more effective to prevent infection with contemporary IAV strains. Reverse genetics methods can be used to introduce mutations into circulating IAV genomes in order to generate novel LAIV (5, 7, 38), including improved MDV (9, 10, 45). We have previously shown that the *ts*, *ca*, and *att* mutations of the A/Ann Arbor/6/60 H2N2 MDV were able to transfer the same phenotype into A/Puerto Rico/8/34 H1N1 (PR8) (10, 51), A/canine/NY/dog23/09 H3N8 (42, 43, 52), and A/equine/Ohio/1/03 H3N8 (46, 47). However, these mutations were not able to transfer the same *ca*, *ts*, and *att* phenotype to the pandemic A/California/04/09 H1N1 (pH1N1) (9, 53), limiting the potential use of this contemporary IAV strain as an MDV for LAIV development (53).

In order to develop a safer MDV based on currently circulating pH1N1, we selected the known *ca*, *ts*, and *att* mutations of the A/Ann Arbor/6/60 H2N2 MDV in combination with the substitution of the conserved leucine at position 319 in PB1 by glutamine (L319Q) (54, 55). We generated recombinant pH1N1 viruses containing either PB1 _L319Q_ alone or in combination with the mutations of the A/Ann Arbor/6/60 H2N2 MDV and evaluated their *in vitro* viral replication phenotypes, as well as their virulence and transmission phenotypes in ferrets and guinea pigs, respectively. Notably, the single amino acid substitution L319Q in the backbone of pH1N1 resulted in a *ts* phenotype *in vitro*, which was associated with reduced levels of virulence and transmission *in vivo*. Moreover, the *ts* and *att* phenotypes observed *in vitro* and *in vivo* were more pronounced when L319Q was combined with the mutations from the A/Ann Arbor/6/60 H2N2 MDV. These results demonstrate the feasibility of generating a safe and attenuated MDV based on the backbone of the contemporary pH1N1, which could be used to update LAIV as a medical countermeasure against currently circulating pH1N1 IAV strains. In addition, this strategy could also be used for the further clinical development of LAIV in <2-year old kids.

## RESULTS

### Generation and characterization of recombinant pH1N1 viruses

To generate the recombinant pH1N1 viruses as candidate MDV, we modified the pH1N1 reverse genetics plasmids to encode the *ts*, *ca*, *att* mutations of the A/Ann Arbor/6/60 H2N2 MDV PB2 (N265S) and PB1 (K391E, E581G, and A661T) and/or the PB1 _L319Q_ mutation (**Figure 1**). No mutation was introduced in the viral NP since pH1N1 NP already encodes a G at position 34. In total we generated six pH1N1 viruses (**Figure 1**), containing different gene constellations: pH1N1 WT (**Figure 1A**), pH1N1 LAIV (**Figure 1B**), pH1N1_PB2 WT/PB1 LAIV_ (**Figure 1C**), pH1N1_PB2 WT/PB1 _L319Q__ (**Figure 1D**), pH1N1_PB2 WT/PB1 LAIV+L319Q_ (**Figure 1E**), and pH1N1_PB2 LAIV/PB1 LAIV+L319Q_ (**Figure 1F**).

**Figure 1.**
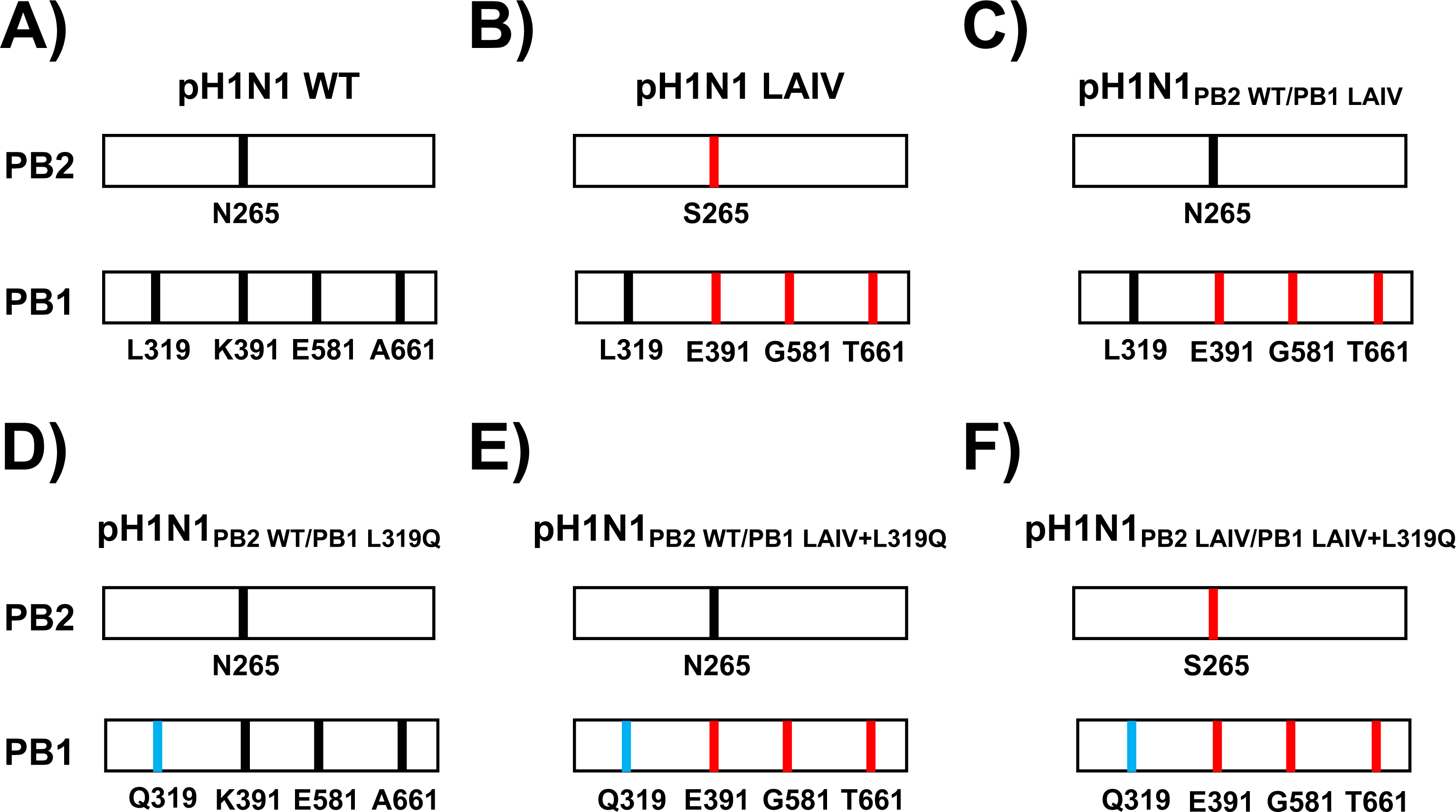
Schematic representation of the PB2 and PB1 viral segments in the different pH1N1 viruses. The PB2 and PB1 viral segments of pH1N1 WT (**A**), pH1N1 LAIV (**B**), pH1N1_PB2 WT/PB1 LAIV_ (**C**), pH1N1_PB2 WT/PB1 _L319Q__ (**D**), pH1N1_PB2 WT/PB1 LAIV+L319Q_ (**E**), and pH1N1_PB2 LAIV/PB1 LAIV+L319Q_ (**F**) viruses are shown. PB2 (N265) and PB1 (L319, K391, E581, and A661) WT amino acid residues are indicated in black. PB2 (S265) and PB1 (E391, G581, and T661) LAIV amino acid changes are indicated in red. PB1 Q319 is shown in blue.

To evaluate the effect of the amino acid changes introduced in the PB2 or PB1 genes on viral polymerase complex activity, a minigenome (MG) assay was performed at three different temperatures (33°C, 37°C, and 39°C) (**Figure 2**). To that end, pDZ ambisense plasmids encoding pH1N1 polymerase complexes (pDZ-PB2WT or PB2LAIV, -PB1 or PB1LAIV or -PB1L319Q or pDZ-PB1LAIV+L319Q, -PA, - and NP), were transiently co-transfected into human 293T cells, together with two viral (v)RNA-like reporter plasmids encoding green fluorescent protein (GFP) and Gaussia luciferase (Gluc) driven by a human RNA polymerase I promoter, and an expression plasmid constitutively expressing Cypridina luciferase (Cluc) under an SV40 promoter (SV40-Cluc), to normalize transfection efficiencies. Then, GFP (**Figure 2A**), and Gluc and Cluc (**Figure 2B**) expression levels were determined at 48 h post-transfection (p.t.) using fluorescent microscopy and luciferase assays, respectively. At 39°C, we observed a significant reduction for both GFP and Gluc expression levels, when a PB1 plasmid containing the L319Q substitutions was included in the polymerase complex. Indeed, MG activity at 39°C was similar for polymerase complexes containing either the full set of LAIV mutations (pH1N1 LAIV) or the single PB1 _L319Q_ mutation (pH1N1_PB2 WT/PB1 L319Q_). When the LAIV and PB1 _L319Q_ mutations were combined, MG activity was greatly reduced at all temperatures (33°C, 37°C), and undetectable at 39°C (pH1N1_PB2 LAIV/PB1 LAIV+L319Q_).

**Figure 2.**
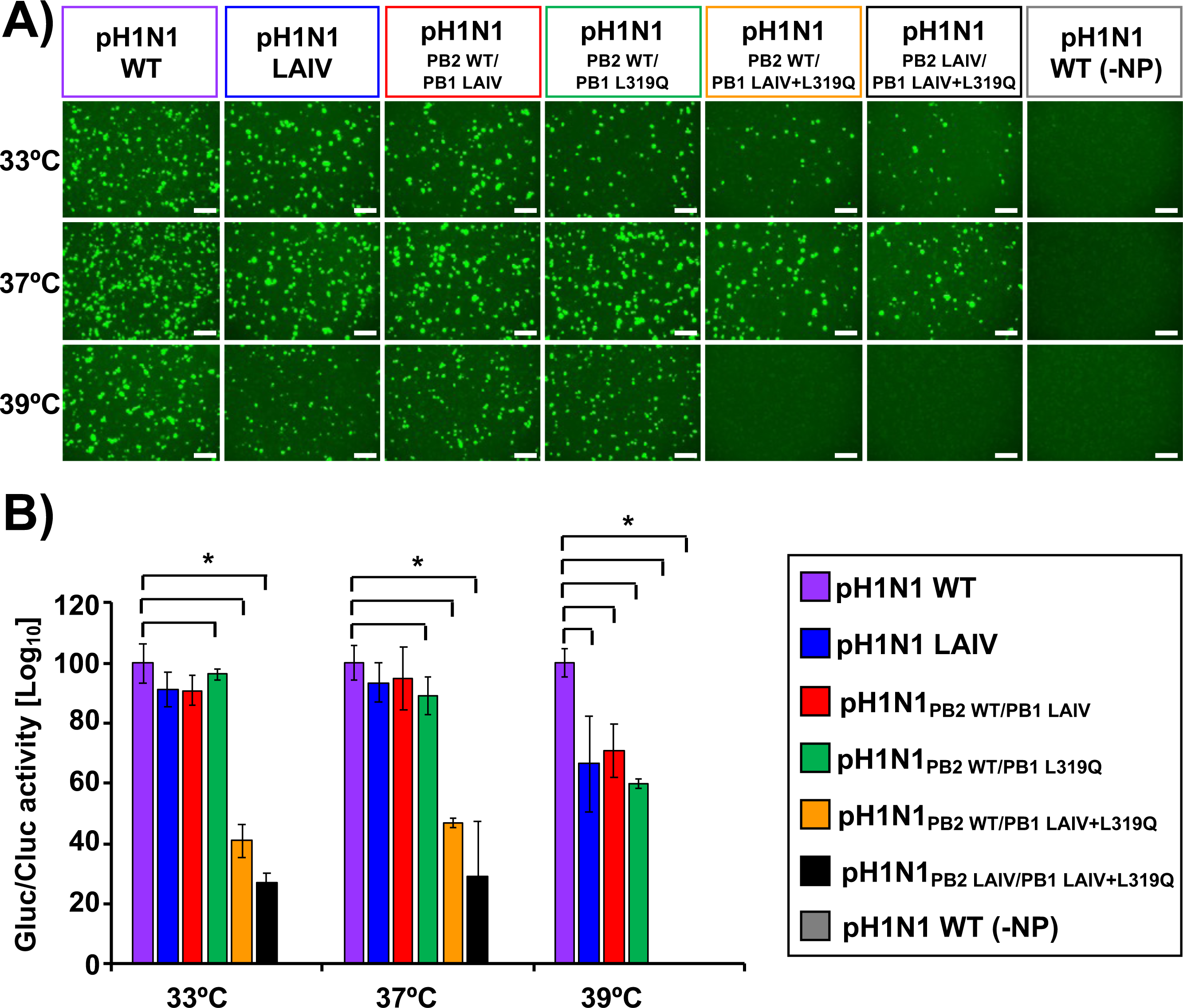
Effect of PB2 and PB1 mutations on pH1N1 polymerase activity at different temperatures. Human 293T cells were transiently co-transfected, using LPF2000, with the indicated combination of ambisense pDZ expression plasmids encoding the minimal requirements for viral genome replication and gene transcription (pDZ-PB2 or PB2_LAIV_, -PB1 or PB1_LAIV_ or -PB1_L319Q_ or pDZ-PB1_LAIV+L319Q_, -PA, and -NP), together with vRNA-like reporter plasmids encoding GFP or Gluc under the control of the human polymerase I promoter and the SV40-Cluc expression plasmid to normalize transfection efficiencies. The MG activity was evaluated at 48 h after transfection by GFP expression under a fluorescence microscope (**A**) or luminescence using a microplate reader (**B**). Gluc activity was normalized to that of Cluc and the data were represented as relative activity considering the activity of pH1N1 WT at each indicated temperature as 100%. Data represent the means and SD of the results determined from triplicate wells. *, P < 0.05 (WT plasmids *versus* other plasmid combinations) using Student’s t test (n = 3 per time point). Scale bar, 100 µm.

To assess whether the introduced mutations from the A/Ann Arbor/6/60 H2N2 MDV conferred a *ts* phenotype and to evaluate the specific contribution of the PB1 _L319Q_ substitution, we performed a multicycle virus growth assay at different temperatures (33°C, 37°C, and 39°C) and compared the results to those for the pH1N1 WT virus (**Figure 3**). For this purpose, MDCK cells were infected at a low multiplicity of infection (MOI 0.001 plaque-forming units (PFU)/cell), and viral titers in cell culture supernatants were determined at 24, 48, 72 or 96 h post-infection (p.i.) (**Figure 3**). Although pH1N1 WT and pH1N1 LAIV replicated to significantly higher titers at 33°C at 24 h p.i., all viruses reached similar levels of replication at the peak of infection (48 to 72 h) (**Figure 3**). However, the virus containing only the PB1 _L319Q_ mutation (pH1N1_PB2 WT/PB1 319Q_) exhibited significantly delayed replication at all temperatures, compared to pH1N1 WT and pH1N1 LAIV (see 24 h p.i. time points; **Figure 3**). Furthermore, viruses containing both the PB1 _L319Q_ mutation and the PB1 _LAIV_ mutations (pH1N1_PB2 WT/PB1 LAIV+L319Q_ and pH1N1_PB2 LAIV/PB1 LAIV+L319Q_) exhibited undetectable levels of replication at all-time points, at both 37°C and 39°C (**Figure 3**). These data show that the PB1 _L319Q_ mutation, alone, is able to confer a *ts* phenotype on pH1N1, and that combining the PB1 _L319Q_ substitution with the mutations of the A/Ann Arbor/6/60 H2N2MDV conferred the strongest *ts* phenotype to pH1N1.

**Figure 3.**
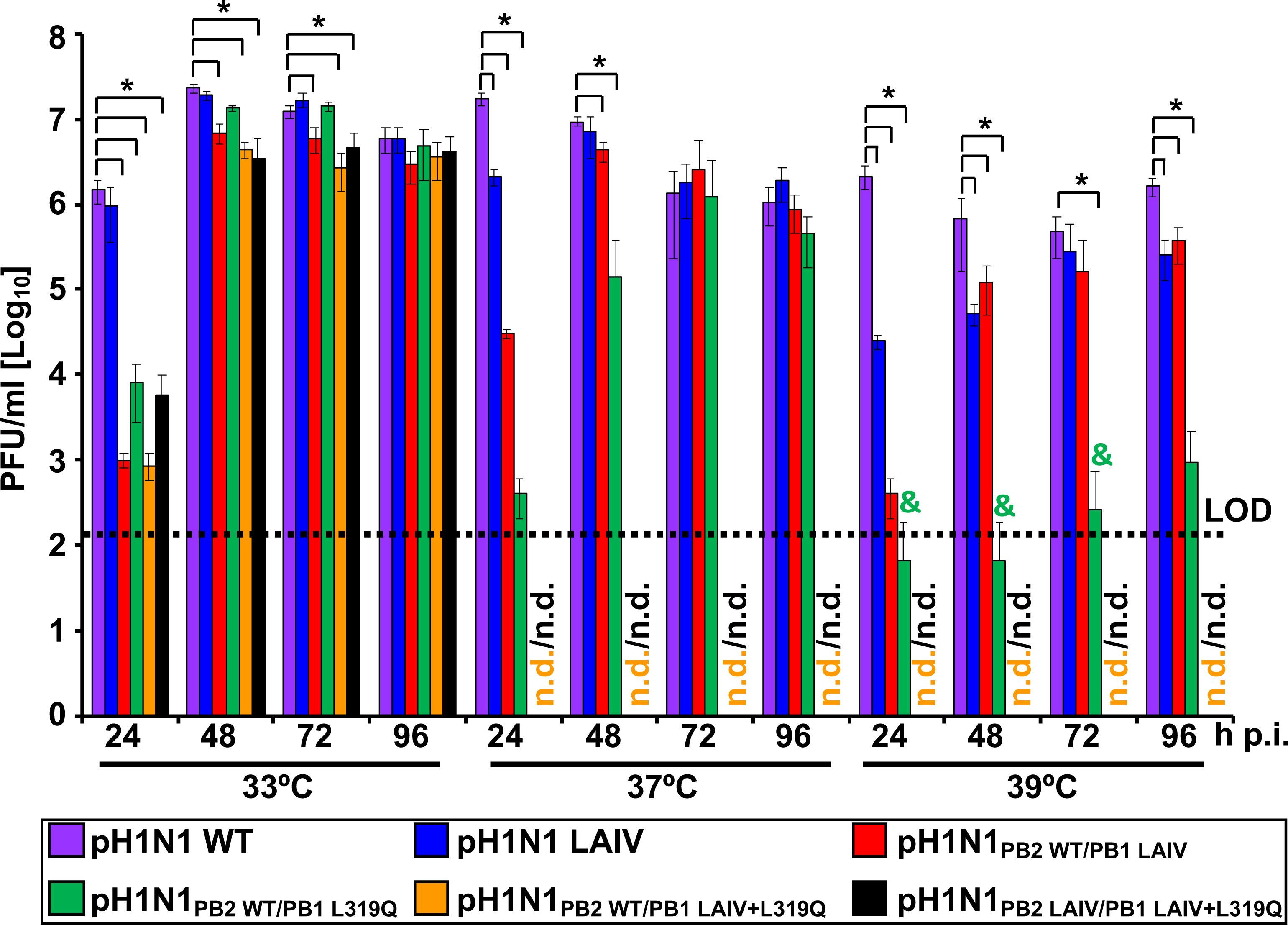
Replication of the pH1N1 viruses in MDCK cells. Cell culture supernatants of MDCK cells infected at a low MOI (0.001) with the indicated WT and mutant viruses at 33°C, 37°C, and 39°C were analyzed at the indicated times p.i. (24, 48, 72 and 96 h) by plaque assay. Data represent the means and SD of the results determined from triplicate wells. The dashed line indicates the limit of detection (LOD) (200 PFU/ml). *, *P* < 0.05 (pH1N1 WT versus pH1N1 mutants) using Student’s *t* test. n.d, non-detected. &,: Virus detected in one of the triplicates.

### Replication and pathogenicity of recombinant pH1N1 viruses in ferrets

Since we observed that the amino acid substitution L319Q had a negative impact in virus replication at higher (37°C or 39°C) temperatures *in vitro* (**Figure 3**), we further assessed the viral replication and pathogenicity of these recombinant pH1N1 viruses in ferrets (**Figure 4**). Groups of ferrets were infected with 10^7^ PFU of each virus, and nasal washes (**Figure 4A**) or oropharyngeal swabs (**Figure 4B**) were collected to examine viral replication on 1 day post-infection (d.p.i.) and 3 d.p.i. In addition, all infected ferrets were euthanized on 4 d.p.i. and virus distribution in the upper respiratory tract (URT) nasal turbinate (**Figure 4C**) and olfactory bulb (**Figure 4D**); and lower respiratory tract (LRT) trachea (**Figure 4E**), and the upper left (UL, **Figure 4F**) or lower left (LL, **Figure 4G**) lungs were also determined. This analysis allowed us to evaluate viral replication and the effect of the A/Ann Arbor/6/60 H2N2 MDV and L319Q mutations, alone or in combination, in a gradient of temperatures, from lower to higher, in the URT or LRT, respectively. Titers of pH1N1 viruses containing the PB1 _L319Q_ mutation (pH1N1_PB2 WT/PB1 L319Q_, pH1N1_PB2 WT/PB1 LAIV+L319Q_ and pH1N1_PB2 LAIV/PB1 LAIV+L319Q_) were consistently lower than those of pH1N1 WT and pH1N1 LAIV in both nasal washes (**Figure 4A**) and oropharyngeal swabs (**Figure 4B**) at 1 d.p.i. However, all viruses displayed similar viral titers at 3 d.p.i. in tissue samples collected at sacrifice (4 d.p.i.), titers of pH1N1 viruses containing the PB1 _L319Q_ mutation were lower than those of pH1N1 WT and pH1N1 LAIV in the LRT (trachea and lung) (**Figures 4E, 4F, and 4G**). However, no significantly differences in viral titers were observed in samples from the URT (**Figures 4C and 4D**). These results suggest that that the PB1 _L319Q_ mutation confers a *ts* and *att* phenotype on pH1N1, which is consistent with our *in vitro* data (**Figures 2 and 3**).

**Figure 4.**
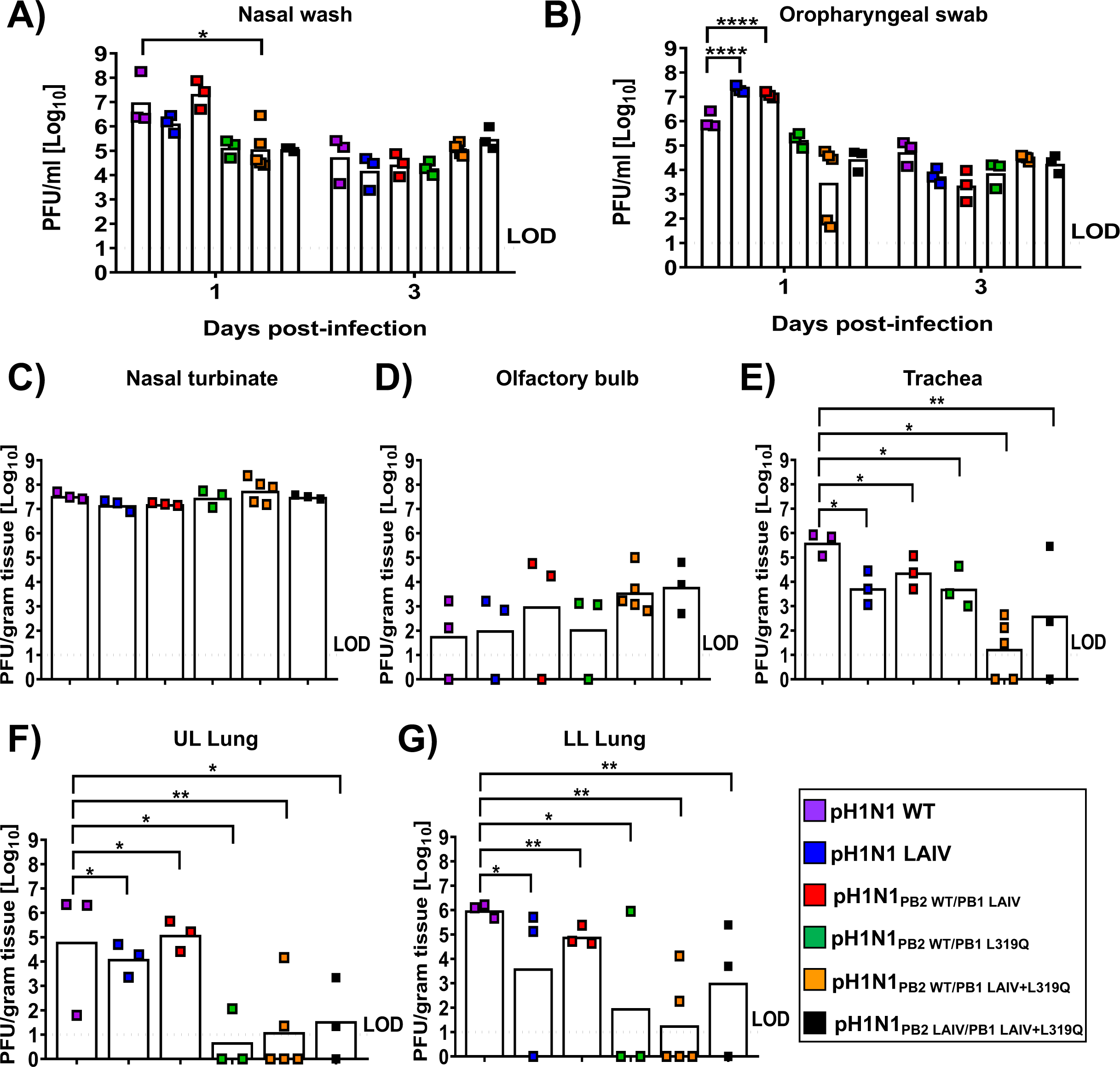
Characterization of the pH1N1 WT and mutant viruses in ferrets. Three to five 4-month-old castrated male Fitch ferrets were inoculated with 10^7^ PFU of the indicated WT and mutant pH1N1 viruses. Viral titers were measured by plaque assay in nasal washes (**A**) and oropharyngeal swabs (**B**) on 1 d.p.i. and 3 d.p.i. as well as in the nasal turbinate (**C**), olfactory bulb (**D**), trachea (**E**); and UL (**F**) or LL (**G**) lungs on day 4 p.i. Dashed lines indicate the limit of detection (LOD; 10 PFU/ml). Data in **A-B** were compared to animals infected with the pH1N1 WT virus and analyzed by two-way ANOVA, followed by a Sidak’s multiple comparison test (multiple time points). Data in **C-G** were compared to pH1N1 WT virus infected animals with one-way ANOVA followed by a Dunnett’s multiple comparison test (single time point). The asterisks refer to the level of significance. *: p<0.05; **: p<0.01; ****: p<0.0001.

In addition to measuring viral titers in tissue homogenates, the upper right lungs of ferrets infected with WT or mutant pH1N1 viruses were also collected at 4 d.p.i., formalin-fixed, embedded, and stained with a pH1N1 polyclonal antibody for immunohistochemical examination of virus distribution and analysis of histopathological changes (**Figure 5A**). The pH1N1 viruses containing the PB1 _L319Q_ substitution (i.e. pH1N1_PB2 WT/PB1 L319Q_, pH1N1_PB2 WT/PB1 LAIV+L319Q_ and pH1N1_PB2 LAIV/PB1 LAIV+L319Q_) showed lower levels of viral replication in the trachea and lung (LRT). Moreover, animals infected with these PB1 _L319Q_ mutant pH1N1 viruses exhibited reduced pathological changes compared to pH1N1 WT or pH1N1 LAIV (e.g., reduced inflammatory infiltrates and necrosis/fibrin deposition), along with lower composite clinical scores (**Figure 5B**). These results echoed our virus replication data of the lungs in **Figure 4**, implying that pH1N1 viruses containing either PB1 _L319Q_ alone or in combination with the mutations of the A/Ann Arbor/6/60 H2N2 MDV acquired attenuated phenotypes in ferrets.

**Figure 5.**
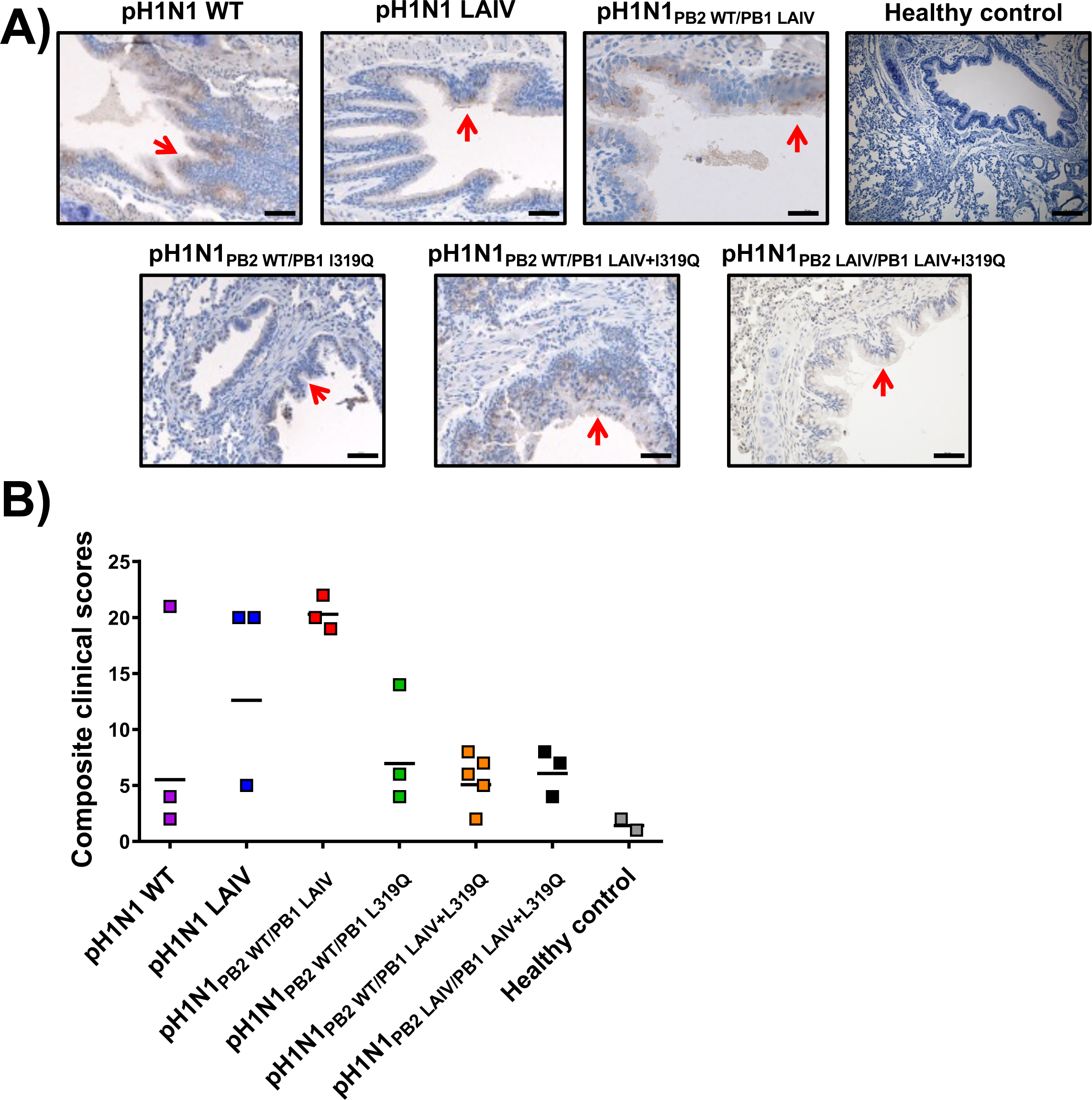
Histopathological analyses of ferrets infected with the WT or LAIV pH1N1 viruses. (**A**) On day 4 p.i., lung tissue from healthy control ferrets or ferrets infected with different pH1N1 viruses in Figure 4 were collected and sections were then analyzed using pH1N1-specific polyclonal antibody and imaged using a Zeiss AxioImager.Z2M. The scale bar denotes 50 µm. Red arrows highlight areas of pH1N1 antigen staining. (**B**) Composite clinical histopathology scores of control or virus-infected ferret lungs were assigned by a blinded veterinary pathologist. Each dot indicates one individual ferret. The black line indicates the geometric mean of the composite clinical scores of each group.

### Replication and transmission efficacy of recombinant pH1N1 viruses in guinea pigs

An important safety criteria for LAIV implementation is that the virus does not transmit from vaccinated individuals to contacts, which are not fit to receive LAIV, such as immunocompromised people. Therefore, we next evaluated the ability of the recombinant pH1N1 viruses to transmit and their ability to prevent pH1N1 WT transmission after vaccination. For these experiments, we used the guinea pig model of IAV transmission (56-58). Because limitations in the number of available animals, the recombinant pH1N1_PB2 LAIV/PB1 LAIV+L319Q_ was not evaluated, given that the levels of replication and attenuation displayed in culture cells and ferrets (**Figures 3 and 4**, respectively) were similar to those of pH1N1_PB2 WT/PB1 LAIV+L319Q_, which was included in the assay. Animals were inoculated intranasally with 10^4^ PFU of pH1N1 WT, pH1N1 LAIV, pH1N1_PB2 WT/PB1 LAIV_, pH1N1_PB2 WT/PB1 L319Q_, and pH1N1_PB2 WT/PB1 LAIV+L319Q_. Then, at 24 h after infection, one naïve guinea pig was exposed to each infected animal by placement in the same cage (56-58). We collected nasal lavage samples from all animals on alternating d.p.i. to follow the kinetics of viral growth in the infected animals and the rate of transmission to exposed cage mates (**Figure 6A**). All pH1N1 viruses replicated in the inoculated guinea pigs (dotted lines), and on 2 d.p.i. we observed increasing viral titers in exposed cage mates (solid lines) for pH1N1 WT, pH1N1 LAIV and pH1N1_PB2 WT/PB1 LAIV_. In contrast, replication of pH1N1_PB2 WT/PB1 LAIV+L319Q_ was delayed in inoculated guinea pigs (**Figure 6F**). Similarly, transmission of viruses containing the PB1 _319Q_ mutation to exposed cage mates was either delayed (pH1N1_PB2 WT/PB1 L319Q_) (**Figure 6E**) or abrogated entirely (pH1N1_PB2 WT/PB1 LAIV+L319Q_) (**Figure 6F**).

**Figure 6.**
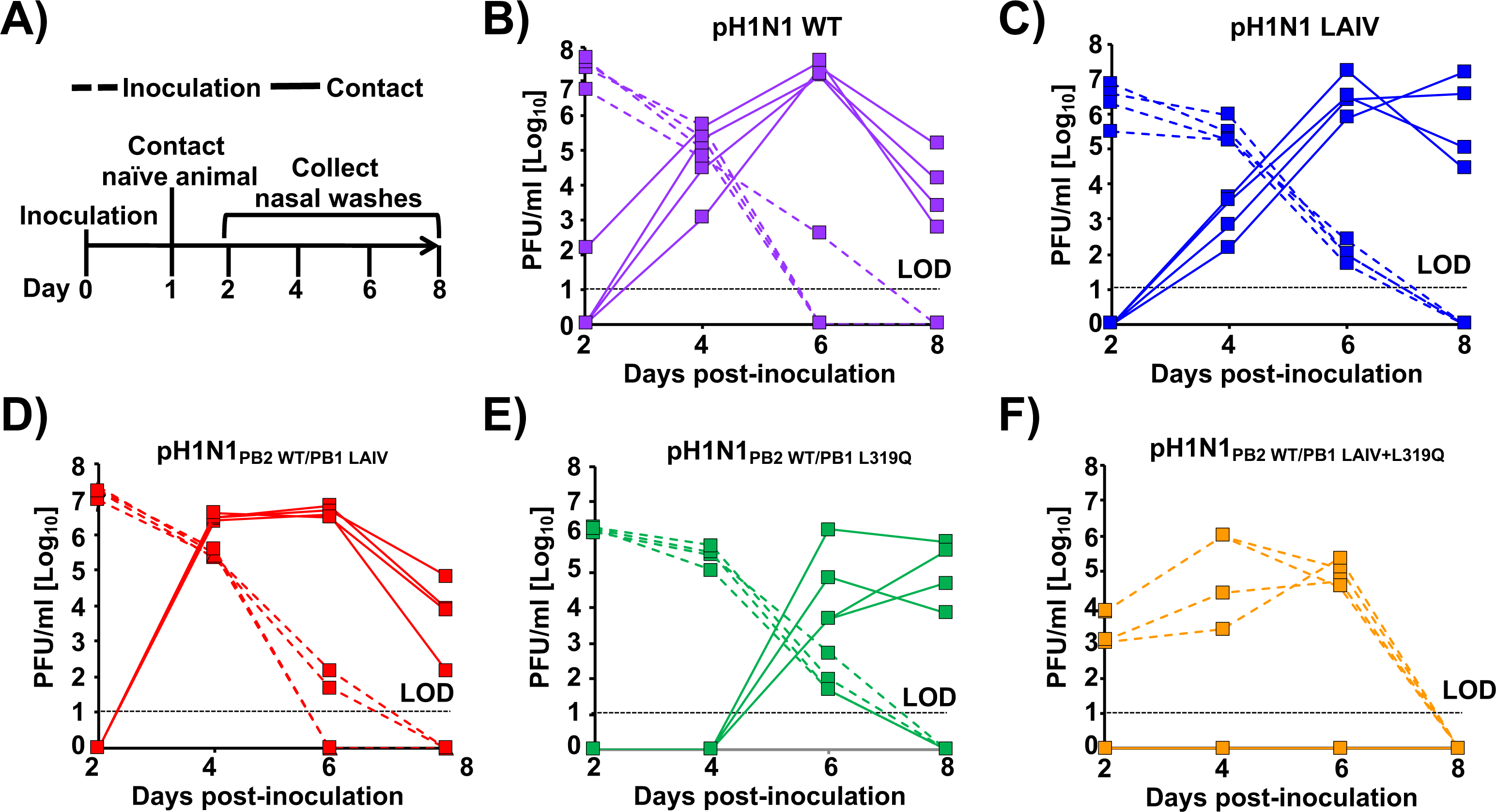
Infection and transmission of WT and mutant pH1N1 viruses in guinea pigs. **A**) Schematic representation of the experimental design: Guinea pigs were infected with 10^4^ PFU of pH1N1 WT (**B**), pH1N1 LAIV (**C**), pH1N1_PB2 WT/PB1 LAIV_ (**D**), pH1N1_PB2 WT/PB1 L319Q_ (**E**), or pH1N1_PB2 WT/PB1 LAIV+L319Q_ (**F**) virus. Then, at 24 h p.i., infected guinea pigs (dashed lines) were placed in a cage with one uninfected guinea pig (solid lines). Nasal washes were collected for 8 days at 48 h intervals, starting at day 2 p.i. for the inoculated animals (24 h post-contact), and virus titers determined by plaque assay (PFU/ml). Dashed horizontal lines indicate the LOD (10 PFU/ml).

Next, we evaluated whether vaccinated guinea pigs were protected against a homologous challenge with pH1N1 WT (**Figure 7**). To that end, animals from the transmission experiment (**Figure 6**; inoculation or contact), were challenged 21 days after initial infection with 10^4^ PFU of pH1N1 WT and viral replication in nasal washes was evaluated for 8 days at 48 h intervals, starting from 2 d.p.i. A group of mock-vaccinated animals was included as a control. All animals positive for the presence of virus in **Figure 6** (both inoculated and contact groups for pH1N1 WT, pH1N1 LAIV, pH1N1_PB2 WT/PB1 LAIV_, and pH1N1_PB2 WT/PB1 L319Q_; and the inoculated group for pH1N1_PB2 WT/PB1 LAIV+L319Q_) were fully protected against pH1N1 WT challenge since virus growth was not detected at any time point analyzed (**Figure 7**). Cage mates (contacts) of animals vaccinated with pH1N1_PB2 WT/PB1 LAIV+L319Q_ were not protected against challenge with pH1N1 WT (**Figure 7B**), as was expected because pH1N1_PB2 WT/PB1 LAIV+L319_ was not transmitted between animals (**Figure 6F**).

**Figure 7.**
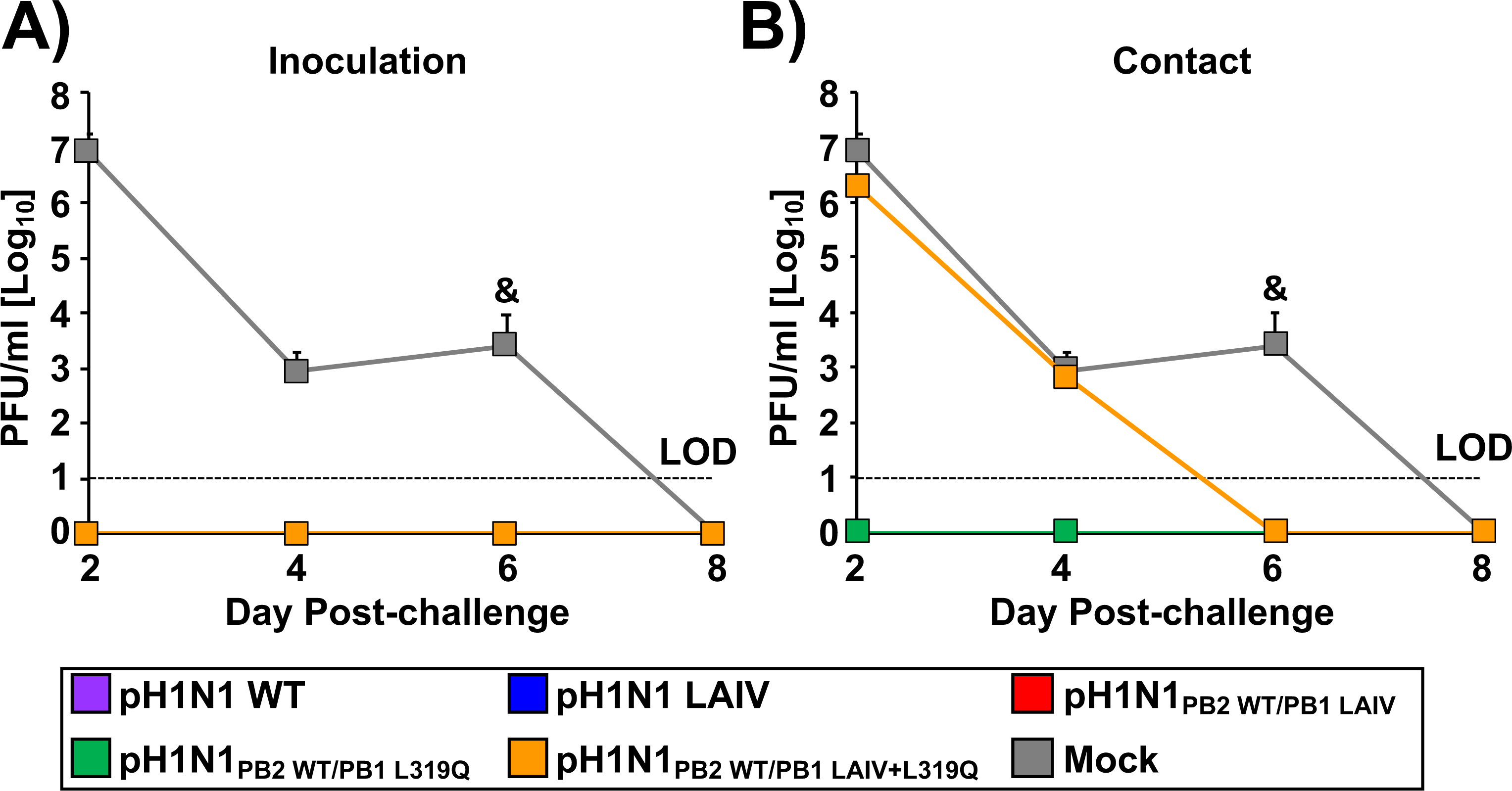
Vaccinated guinea pigs are protected against homologous challenge with pH1N1 WT. Guinea pigs from the transmission experiment shown in Figure 6 were challenged 21 days later with 10^4^ PFU of pH1N1 WT. One group of mock-vaccinated animals was included as a control. To evaluate viral replication, nasal washes were collected for 8 days at 48 h intervals, starting from 2 day p.i., and virus titers were determined by plaque assay (PFU/ml). Dashed lines indicate the LOD (10 PFU/ml). &,: Virus detected in one out of the 4 guinea pigs. The same mock control group has been included in both panels A and B.

## DISCUSSION

A single amino acid change in PB2 (N265S), three substitutions in PB1 (K391E, E581G and A661T) and a mutation in NP (D34G) account for the *ts*, *ca* and *att* phenotype of the current U.S. A/Ann Arbor/6/60 H2N2 MDV (5, 7, 9, 10, 38, 48-50). In addition, we and others have shown that the amino acid substitutions responsible for the *ts*, *ca*, and *att* of the A/Ann Arbor/6/60 H2N2 MDV can confer the same phenotype to other IAV strains (10, 42, 43, 46, 47, 51, 52). However, these mutations fail to attenuate strains of IAV such as some avian influenza viruses and the pH1N1 (9, 38, 53, 59). In order to overcome this reduced attenuation, it may be necessary to introduce additional mutations into the genome background of pH1N1 to generate an updated MDV suitable for use in LAIV capable of protecting against contemporary circulating H1N1 strains. For instance, we have shown that the *ts*, *ca*, *att* mutations from the Russian A/Leningrad/17/57 H2N2 MDV confer a greater *ts*, and *att* phenotype to pH1N1 than the mutations of the U.S. A/Ann Arbor/6/60 H2N2 MDV (9).

In this study, we replaced the conserved leucine at PB1 residue 319 with glutamine (55) into pH1N1, either alone or in combination, with the other A/Ann Arbor/6/60 H2N2 MDV mutations in PB2 and PB1 (**Figure 1**). First, the effect of the A/Ann Arbor/6/60 H2N2 MDV and/or PB1 _L319Q_ mutations on viral polymerase activity was evaluated using a MG assay (**Figure 2**). We observed a reduction in polymerase activity when a PB1 plasmid containing the LAIV+L319Q substitutions was incorporated in the polymerase complex, and this reduction was more significant at higher temperatures. Next, we generated recombinant pH1N1 viruses containing different PB2 and PB1 combinations (pH1N1 WT, pH1N1 LAIV, pH1N1_PB2 WT/PB1 LAIV_, pH1N1_PB2 WT/PB1 L319Q_, pH1N1_PB2 WT/PB1 LAIV+L319Q_, and pH1N1_PB2 LAIV/PB1 LAIV+L319Q_) and evaluated their replication *in vitro* (**Figure 3**) and in two relevant animal models for IAV infection and transmission: ferrets (**Figures 4 and 5**) and guinea pigs (**Figures 6** **and 7**), respectively.

The PB1 _L319Q_ substitution, alone, was sufficient to increase the *ts* phenotype of pH1N1 in culture cells in both MG (**Figure 2**) and virus replication assays (**Figure 3**). Moreover, viruses containing PB1 _L319Q_ were also attenuated in ferrets (**Figures 4 and 5**), and exhibited reduced transmission in guinea pigs (**Figures 6** **and 7**). Notably, we observed an enhancement of the *ts*, and *att* phenotypes in pH1N1 when we combined the A/Ann Arbor/6/60 H2N2 MDV mutations with the PB1 _L319Q_ substitution. Consistent with this, the replication of pH1N1 viruses containing the PB1 _L319Q_ amino acid change was reduced in the LRT of ferrets (**Figure 4**), suggesting that the PB1 _L319Q_ substitution, alone, can confer an *att* phenotype in this pH1N1 strain (and that it can increase the *att* phenotype of pH1N1 LAIV). Moreover, our guinea pig data showed that pH1N1_PB2 WT/PB1 LAIV+L319Q_ was not detectably transmitted from experimentally-infected animals to naive cage mates (**Figure 6**).

In order to understand better the basis for the observed effect of the PB1 _L319Q_ substitution, we examined available crystal structures of the IAV RNA-dependent RNA polymerase (RdRp) complex. For this purpose, we used the recently reported IAV polymerase heterotrimer structure for A/Northern Territory/60/68 H3N2 (PDB ID 6RR7) (60) as a reference model, since it possesses >97% amino acid identity with the pH1N1 virus polymerase (PB2: 97%, PB1: 98% and PA: 97%). We analyzed the location of the A/Ann Arbor/6/60 H2N2 MDV mutations, and the PB1 _L319Q_ substitution (**Figure 8**). We observed that amino acid residue 319 in PB1 is located near the interface between the PB1 and PA subunits, and in close physical proximity to PB1 residue 391 (a mutation present in the A/Ann Arbor/6/60 H2N2 MDV). Therefore, the amino acid change PB1 _L319Q_ could affect polymerase complex formation and/or stability. Further studies will be necessary to test this hypothesis, and to gain a better mechanistic understanding of how PB1 _L319Q_ enhances the *ts*, *att* phenotypes of pH1N1 containing the A/Ann Arbor/6/60 H2N2 MDV mutations. However, a recent study has suggested that a PR8 virus containing PB1 _L319Q_ and PB2 N265S amino acid changes has propensity to generate high levels of semi-infectious particles at 39°C, while this did not occur at 33°C (54). In addition, viruses containing the LAIV mutations alone also demonstrated this same effect (54). Chen et al. has also showed that the attenuating mutations of LAIV altered IAV M1 protein levels and virion morphology in a temperature sensitive manner (61).

**Figure 8.**
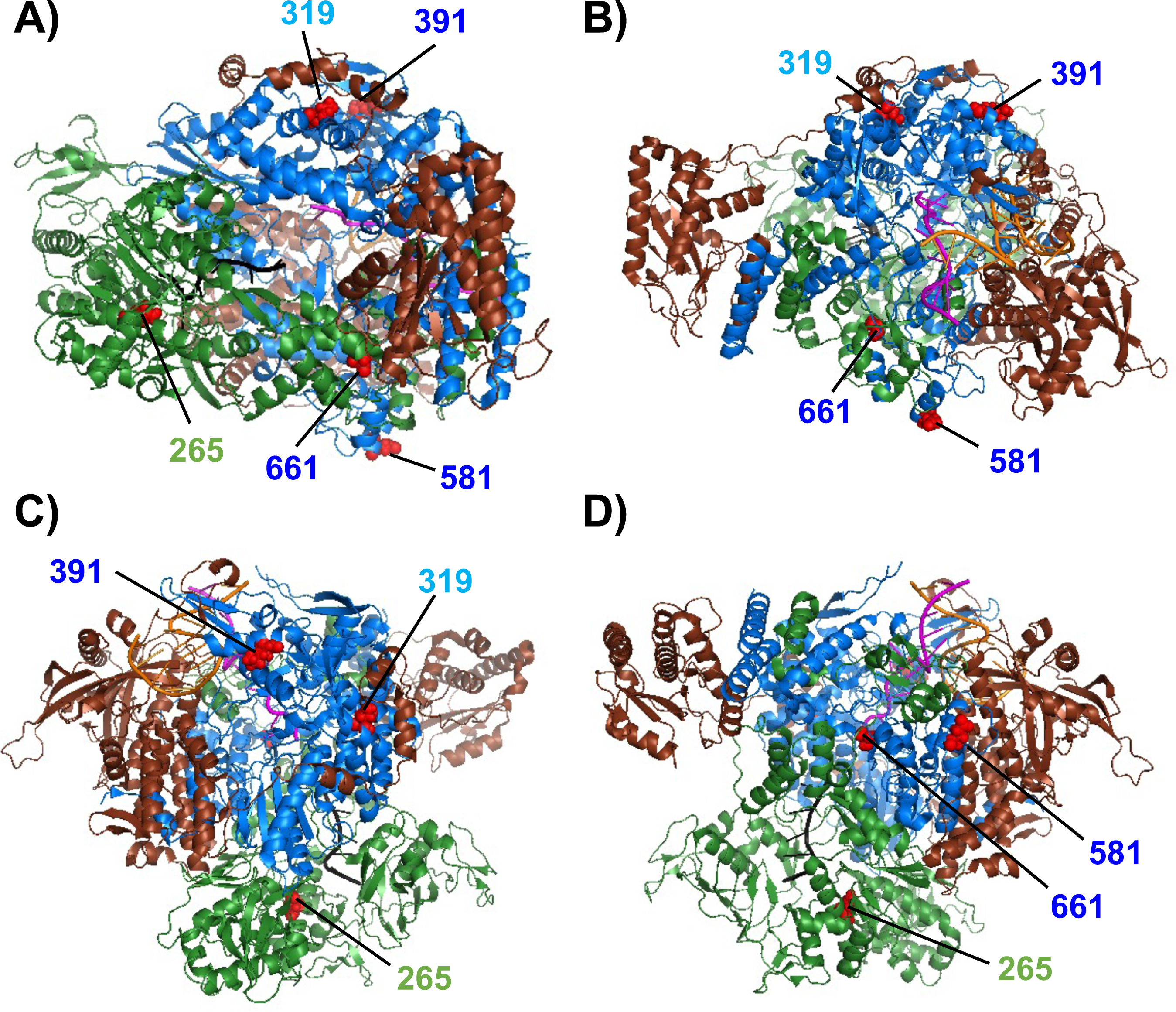
Structure of the IAV polymerase complex. The IAV polymerase heterotrimer complex bound to 3‘ and 5’ vRNA promoters and capped RNA primer complex (PDB code 6RR7) is shown in four different views (**A to D**) using the same coloring code: PA (brown), PB1 (blue), and PB2 (green). The 3‘ and 5’ vRNA promoters, and capped RNA primer, are colored in orange, magenta and black, respectively. The locations of the A/AnnArbor/6/60 H2N2 MDV PB2 265 (green) and PB1 391, 581, and 661 (dark blue) mutations; and the PB1 319 residue (light blue) are shown on the background of the polymerase complex colored in red.

Collectively, the results reported here demonstrate that the mutations responsible for the *ts*, *ca*, and *att* phenotypes of the A/Ann Arbor/6/60 H2N2 MDV fail to confer the same phenotype on pH1N1. However, by combining them (in whole or in part) with the PB1 _L319Q_ mutation, they can confer robust *att* and *ts* phenotypes on pH1N1. Moreover, PB1 _L319Q_ alone can confer a substantial level of *in vivo* attenuation on pH1N1 virus. Future studies will determine whether the PB1 _L319Q_ mutation can also attenuate other IAV strains, including avian influenza viruses that are not attenuated by the mutations of the A/Ann Arbor/6/60 H2N2 MDV (9, 38, 53, 59). Finally, our data support the feasibility of using PB1 _L319Q_ to attenuate contemporary virus backbones as the basis for creating an updated MDV that may improve the efficacy and safety of LAIV, by better matching the vaccine to currently circulating virus strains. Vaccine immunogenicity and safety are determined by characteristics of both the vaccine and the host. For instance, current LAIV are contraindicated for most people who are severely immunocompromised. Therefore, there is a necessity to develop alternative LAIV with better safety profiles to target population that not fit requirements to receive current LAIV. So, PB1 _L319Q_ amino acid change could be combined with other mutations to generate more attenuated viruses, but still able to induce potent humoral and cellular responses against circulating IAV strains. Such strategy could also facilitate the clinical development of LAIV for the <2-year old kids.

## MATERIAL AND METHODS

### Cells and viruses

Human embryonic kidney 293T (293T; ATCC CRL-11268) and Madin-Darby canine kidney (MDCK; ATCC CCL-34) cells were grown at 37°C (and air enriched with 5% CO2) in Dulbecco’s modified Eagle’s medium (DMEM; Mediatech, Inc.) supplemented with 5% fetal bovine serum (FBS; Atlanta Biologicals) and 1% penicillin (100 units/ml)– streptomycin (100 μg/ml)–2 mM L-glutamine (P-S-G; Mediatech, Inc.). Recombinant wild-type (WT) and LAIV pandemic A/California/4_NYICE_E3/2009 H1N1 (pH1N1) have been previously described (53, 62-66).

### Rescue of recombinant pH1N1 viruses

Ambisense pDZ-PB2_LAIV_ and pDZ-PB1_LAIV_ plasmids containing the temperature sensitive (*ts*), cold-adapted (c*a*), and attenuated (*att*) mutations of the A/Ann Arbor/6/60 H2N2 MDV in PB2 (N265S) and PB1 (K391E, D581G, and A661T), respectively, were previously described (53). To generate the recombinant pH1N1 viruses containing PB1 _L319Q_, ambisense plasmids pDZ-PB1L319Q or pDZ-PB1LAIV+L319Q were generated using standard molecular cloning methods. Recombinant pH1N1 viruses were recovered as previously (67-69). Briefly, co-cultures (1:1) of human 293T and MDCK cells in 6-well plates were co-transfected in suspension with 1 µg of each of the ambisense plasmids (pDZ-PB2 or PB2_LAIV_, -PB1 or PB1LAIV or -PB1L319Q or pDZ-PB1LAIV+L319Q, -PA, -HA, -NP,-NA, -M and -NS) using Lipofectamine 2000 (Invitrogen). At 12 h p.t., transfection medium was replaced with DMEM containing 0.3% bovine serum albumin (BSA), 1% P-S-G, and 0.5 µg/ml of N-tosyl-L-phenylalanine chloromethyl ketone (TPCK)-treated trypsin (Sigma). At 72 h p.t., cell culture supernatants were collected, clarified, and used to infect fresh monolayers of MDCK cells (10^6^ cells/well, 6-well plate format). At 3-4 d.p.i., recombinant viruses were plaque purified and scaled up in MDCK cells (67). Virus stocks were produced at 33°C and titrated by plaque assay (PFU/ml) on MDCK cells (67, 68). Virus stocks were confirmed by sequencing (ACGT Inc.) the PB2 and PB1 open reading frames (ORFs) using purified total RNA (Trizol reagent, Invitrogen) from infected MDCK cells (10^6^ cells/well, 6-well plate format).

### Minigenome (MG) assays

To evaluate the effect of WT and mutant PB2 and PB1 proteins on viral polymerase activity, a MG assay was performed as previously described (42, 53). Briefly, human 293T cells (5 × 10^5^ cells/well, 12-well plate format, triplicates) were transiently co-transfected in suspension, using Lipofectamine 2000 (Invitrogen), with 125 ng of each ambisense pDZ plasmids (pDZ-PB2 or PB2_LAIV_, -PB1 or PB1_LAIV_ or -PB1_L319Q_ or pDZ-PB1_LAIV+L319Q_, - PA, - and NP), together with 250 ng of two reporter viral (v)RNA-like expression pPOL-I plasmids encoding GFP or Gluc driven by a human RNA polymerase I promoter (53, 70). A Cluc-encoding plasmid under the simian virus 40 promoter (SV40-Cluc, 50 ng) was included to normalize transfection efficiencies (71, 72). Cells transfected in the absence of pDZ NP were used as a negative control. At 6 h p.t., cells were placed at 33°C, 37°C, or 39°C, and viral replication and transcription were evaluated at 48 h p.t. GFP was imaged using a fluorescence microscope, while Gluc and Cluc expression levels were determined using Biolux Gaussia or Cypridina luciferase assay kits (New England BioLabs) and a microplate reader. The mean value and standard deviation (SD) were calculated using Microsoft Excel software.

### Virus growth kinetics

To evaluate virus replication properties in tissue culture cells, confluent monolayers of MDCK cells (4x10^5^ cells/well, 12-well plate format, triplicates) were infected at a MOI of 0.001 PFU/cell. After 1 h of virus adsorption at room temperature, cells were overlaid with DMEM containing 0.3% BSA, 1% P-S-G, and 1 µg/ml of TPCK-treated trypsin. Cells were incubated at 33°C, 37°C, or 39°C, and cell culture supernatants were collected at the indicated times (24, 48, 72 and 96 h.p.i.) to determine the viral titers by standard plaque assay (PFU/ml) as described below.

### Plaque assays

Confluent monolayers of MDCK cells (10^6^ cells/well, 6-well plate format, triplicates) were infected as indicated above and after 1 h virus adsorption at room temperature, cells were overlaid with agar-containing culture medium and incubated at 33°C for 3 days. Then, cells were fixed with 4% formaldehyde, the overlays were removed, and cells were stained with crystal violet (67, 68).

### Ferret immunization

All ferret experiments were performed in accordance with an animal protocol (IACUC-2017-0136) approved by the Institutional Animal Care and Use Committee (IACUC) of the Icahn School of Medicine at Mount Sinai (ISMMS) All ferrets were housed in a temperature- and relative humidity-controlled Animal Biosafety Level 2 (ABSL-2) facility. All procedures were performed as previously described (73). Briefly, four-month-old influenza virus-seronegative outbred castrated male Fitch ferrets were purchased from Triple F Farms (Gillett, PA, USA). Animals were randomly assigned to the infection or control groups. Groups of ferrets (n=3 for each group, except for group inoculated with pH1N1_PB2 WT/PB1 LAIV+L319Q_, where 5 animals were used) were infected with either pH1N1 WT virus or pH1N1 LAIV groups (pH1N1 LAIV, pH1N1_PB2 WT/PB1 LAIV_, pH1N1_PB2 WT/PB1 L319Q_, pH1N1_PB2 WT/PB1 LAIV+L319Q_, or pH1N1_PB2 LAIV/PB1 LAIV+L319Q_) at a dosage of 10^7^ PFU per animal. A PBS control group was included as a negative control. To examine the pathogenicity of the WT or LAIV pH1N1 viruses, nasal wash and oropharyngeal swab samples were collected from anesthetized ferrets on 1 d.p.i. and 3 d.p.i. On 4 d.p.i., ferrets were euthanized by exsanguination followed by intracardiac injection of Sleepaway euthanasia solution (Fort Dodge, Sodium Pentobarbital). Tissue specimens (olfactory bulbs, nasal turbinates, trachea, upper left (UL), and lower left (LL) lungs) were collected from each individual ferret to quantify viral titers by plaque assay. The upper right lung of each ferret was collected, perfused, and fixed with 10% Formalin for immunohistochemistry (IHC) stain.

### Ferret histopathology evaluation

Histopathology and blinded scoring were performed by the Comparative Pathology Laboratory (CPL) at ISMMS. Formalin-fixed, paraffin-embedded (FFPE) lung specimens obtained from pH1N1 WT or pH1N1 LAIV (pH1N1 LAIV, pH1N1_PB2 WT/PB1 LAIV_, pH1N1_PB2 WT/PB1 L319Q_, pH1N1_PB2 WT/PB1 LAIV+L319Q_, or pH1N1_PB2 LAIV/PB1 LAIV+L319Q_)-infected, or mock-treated ferrets were cut into 5 μm sections and stained with hematoxylin and eosin (H&E) for histopathological assessment. In addition, we used an A/California/04/09 H1N1 polyclonal antibody for IHC stain to examine the virus distributions in the lung. All sections were evaluated by a veterinary pathologist who was blinded to the infection groups. According to the extent of epithelial degeneration, damage, and/or necrosis, and the level of inflammation, it was scored as 0 (None), 1 (Mild), 2 (Moderate), 3 (Marked), and 4 (Severe). Transmitted-light brightfield images were captured by a Zeiss Axio Imager.Z2 microscope and processed with the ZEISS ZENpro microscope software.

### Guinea pig immunization

All work with guinea pigs was approved by the Emory Institutional Animal Care and Use Committee (protocol number PROTO201700595). Female Hartley strain guinea pigs were obtained from Charles River Laboratories. Intranasal inoculation and nasal lavage were performed as described (56-58). Prior to intranasal inoculation and nasal lavage, the guinea pigs were sedated with a mixture of ketamine and xylazine (30 mg/kg of body weight and 4 mg/kg, respectively). Animals were inoculated (n = 4) with 10^4^ PFU of the indicated viruses diluted in PBS. Following inoculation and recovery from sedation, animals were housed in Caron 6040 environmental chambers set to 5°C and 20% relative humidity. At 24 h p.i. of the donor animals, contact guinea pigs were introduced into the same cage with the donor animals. Virus replication was evaluated by determining viral titers in the nasal washes collected at 2, 4, 6, and 8 d.p.i. by plaque assay as described above.

### Statistical Analysis

Microsoft Excel (Microsoft Corporation) was necessary to perform some of the calculations, and to visualize the raw data. GraphPad Prism software (v8.0.1) was used to perform the indicated statistical analysis.

## ACKNOWLEDGMENTS

This research was partially funded by the New York Influenza Center of Excellence (NYICE) (HHSN 272201400005C), and Emory-UGA (HHSN 272201400004C), members of the National Institute of Allergy and Infectious Diseases (NIAID), National Institutes of Health (NIH), Department of Health and Human Services, Centers of Excellence for Influenza Research and Surveillance (CEIRS) network. The work was also supported by grants W81XWH-18-1-0460-PRMRP-DA (to L.M.-S.) and W81XWH-17-1-0168 (to S.D.) from the Department of Defense (DoD) Peer Reviewed Medical Research Program (PRMRP), as well as NIH R01 AI145332-01 (to L.M.-S.). This research was also partly funded by NIAID grant P01AI097092, by CEIRR (Center for Research on Influenza Pathogenesis and Transmission), a NIAID funded Center of Excellence for Influenza Research and Response (CEIRR, contract # 75N93021C00014) and by the Collaborative Influenza Vaccine Innovation Centers NIAID contract 75N93019C00051 to AG-S. A.N. received a “Ramon y Cajal” Incorporation grant (RYC-2017) from the Spanish Ministry of Science, Innovation. Finally, A.C. received support from NIH grants T32GM068411 and T32GM007356.

## REFERENCES

1. Bouvier NM, Palese P. 2008. The biology of influenza viruses. Vaccine 26 Suppl 4:D49–53.

2. Shaw MP, P. 2007. Orthomyxoviridae: the viruses and their replication. In: Knipe, DM.; Howley, PM.; Griffin, DE.; Lamb, RA.; Martin, MA., editors. Fields Virology 5th Lippincott Williams and WIlkins: PA, USA.

3. Wille M, Holmes EC. 2019. The Ecology and Evolution of Influenza Viruses. Cold Spring Harb Perspect Med.

4. Tesini BL, Kanagaiah P, Wang J, Hahn M, Halliley JL, Chaves FA, Nguyen PQT, Nogales A, DeDiego ML, Anderson CS, Ellebedy AH, Strohmeier S, Krammer F, Yang H, Bandyopadhyay S, Ahmed R, Treanor JJ, Martinez-Sobrido L, Golding H, Khurana S, Zand MS, Topham DJ, Sangster MY. 2019. Broad Hemagglutinin-Specific Memory B Cell Expansion by Seasonal Influenza Virus Infection Reflects Early-Life Imprinting and Adaptation to the Infecting Virus. J Virol 93.

5. Blanco-Lobo P, Nogales A, Rodriguez L, Martinez-Sobrido L. 2019. Novel Approaches for The Development of Live Attenuated Influenza Vaccines. Viruses 11.

6. Clark AM, DeDiego ML, Anderson CS, Wang J, Yang H, Nogales A, Martinez-Sobrido L, Zand MS, Sangster MY, Topham DJ. 2017. Antigenicity of the 2015-2016 seasonal H1N1 human influenza virus HA and NA proteins. PLoS One 12:e0188267.

7. Nogales A, Martinez-Sobrido L. 2016. Reverse Genetics Approaches for the Development of Influenza Vaccines. Int J Mol Sci 18.

8. Topham DJ, DeDiego ML, Nogales A, Sangster MY, Sant A. 2019. Immunity to Influenza Infection in Humans. Cold Spring Harb Perspect Med.

9. Smith A, Rodriguez L, Ghouayel ME, Nogales A, Chamberlain JM, Sortino K, Reilly E, Feng C, Topham DJ, Martinez-Sobrido L, Dewhurst S. 2019. A live-attenuated influenza vaccine (LAIV) elicits enhanced heterologous protection when the internal genes of the vaccine are matched to the challenge virus. J Virol.

10. Rodriguez L, Blanco-Lobo P, Reilly EC, Maehigashi T, Nogales A, Smith A, Topham DJ, Dewhurst S, Kim B, Martinez-Sobrido L. 2019. Comparative Study of the Temperature Sensitive, Cold Adapted and Attenuated Mutations Present in the Master Donor Viruses of the Two Commercial Human Live Attenuated Influenza Vaccines. Viruses 11.

11. Rajao DS, Perez DR. 2018. Universal Vaccines and Vaccine Platforms to Protect against Influenza Viruses in Humans and Agriculture. Front Microbiol 9:123.

12. Mameli C, D’Auria E, Erba P, Nannini P, Zuccotti GV. 2018. Influenza vaccine response: future perspectives. Expert Opin Biol Ther 18:1–5.

13. Grohskopf LA, Sokolow LZ, Broder KR, Olsen SJ, Karron RA, Jernigan DB, Bresee JS. 2016. Prevention and Control of Seasonal Influenza with Vaccines. MMWR Recomm Rep 65:1–54.

14. Girard MP, Cherian T, Pervikov Y, Kieny MP. 2005. A review of vaccine research and development: human acute respiratory infections. Vaccine 23:5708–24.

15. Chung JR, Flannery B, Thompson MG, Gaglani M, Jackson ML, Monto AS, Nowalk MP, Talbot HK, Treanor JJ, Belongia EA, Murthy K, Jackson LA, Petrie JG, Zimmerman RK, Griffin MR, McLean HQ, Fry AM. 2016. Seasonal Effectiveness of Live Attenuated and Inactivated Influenza Vaccine. Pediatrics 137:e20153279.

16. MG Thompson DS, H Zhou, CB Bridges, PY Cheng, E Burns, JS Bresee, NJ Cox, Influenza Div, National Center for Immunization and Respiratory Diseases, CDC. 2010. Updated Estimates of Mortality Associated with Seasonal Influenza through the 2006-2007 Influenza Season. http://www.ncbi.nlm.nih.gov/pubmed/22895385.

17. Johnson KEE, Ghedin E. 2019. Quantifying between-Host Transmission in Influenza Virus Infections. Cold Spring Harb Perspect Med.

18. Sutton TC. 2018. The Pandemic Threat of Emerging H5 and H7 Avian Influenza Viruses. Viruses 10.

19. Taubenberger JK, Morens DM. 2019. The 1918 Influenza Pandemic and Its Legacy. Cold Spring Harb Perspect Med.

20. Chen R, Holmes EC. 2008. The evolutionary dynamics of human influenza B virus. J Mol Evol 66:655–63.

21. Garten RJ, Davis CT, Russell CA, Shu B, Lindstrom S, Balish A, Sessions WM, Xu X, Skepner E, Deyde V, Okomo-Adhiambo M, Gubareva L, Barnes J, Smith CB, Emery SL, Hillman MJ, Rivailler P, Smagala J, de Graaf M, Burke DF, Fouchier RA, Pappas C, Alpuche-Aranda CM, Lopez-Gatell H, Olivera H, Lopez I, Myers CA, Faix D, Blair PJ, Yu C, Keene KM, Dotson PD, Jr., Boxrud D, Sambol AR, Abid SH, St George K, Bannerman T, Moore AL, Stringer DJ, Blevins P, Demmler-Harrison GJ, Ginsberg M, Kriner P, Waterman S, Smole S, Guevara HF, Belongia EA, Clark PA, Beatrice ST, Donis R, et al. 2009. Antigenic and genetic characteristics of swine-origin 2009 A(H1N1) influenza viruses circulating in humans. Science 325:197–201.

22. Long JS, Mistry B, Haslam SM, Barclay WS. 2019. Host and viral determinants of influenza A virus species specificity. Nat Rev Microbiol 17:67–81.

23. Parrish CR, Murcia PR, Holmes EC. 2015. Influenza virus reservoirs and intermediate hosts: dogs, horses, and new possibilities for influenza virus exposure of humans. J Virol 89:2990–4.

24. Xu X, Lindstrom SE, Shaw MW, Smith CB, Hall HE, Mungall BA, Subbarao K, Cox NJ, Klimov A. 2004. Reassortment and evolution of current human influenza A and B viruses. Virus Res 103:55–60.

25. Korenkov D, Isakova-Sivak I, Rudenko L. 2018. Basics of CD8 T-cell immune responses after influenza infection and vaccination with inactivated or live attenuated influenza vaccine. Expert Rev Vaccines 17:977–987.

26. Schotsaert M, Garcia-Sastre A. 2017. Inactivated influenza virus vaccines: the future of TIV and QIV. Curr Opin Virol 23:102–106.

27. McLean HQ, Caspard H, Griffin MR, Poehling KA, Gaglani M, Belongia EA, Talbot HK, Peters TR, Murthy K, Ambrose CS. 2017. Effectiveness of live attenuated influenza vaccine and inactivated influenza vaccine in children during the 2014-2015 season. Vaccine 35:2685–2693.

28. Hoft DF, Lottenbach KR, Blazevic A, Turan A, Blevins TP, Pacatte TP, Yu Y, Mitchell MC, Hoft SG, Belshe RB. 2017. Comparisons of the Humoral and Cellular Immune Responses Induced by Live Attenuated Influenza Vaccine and Inactivated Influenza Vaccine in Adults. Clin Vaccine Immunol 24.

29. Del Giudice G, Rappuoli R. 2015. Inactivated and adjuvanted influenza vaccines. Curr Top Microbiol Immunol 386:151–80.

30. Cheng X, Zengel JR, Suguitan AL, Jr., Xu Q, Wang W, Lin J, Jin H. 2013. Evaluation of the humoral and cellular immune responses elicited by the live attenuated and inactivated influenza vaccines and their roles in heterologous protection in ferrets. J Infect Dis 208:594–602.

31. Belshe RB, Edwards KM, Vesikari T, Black SV, Walker RE, Hultquist M, Kemble G, Connor EM, Group C-TCES. 2007. Live attenuated versus inactivated influenza vaccine in infants and young children. N Engl J Med 356:685–96.

32. Basha S, Hazenfeld S, Brady RC, Subbramanian RA. 2011. Comparison of antibody and T-cell responses elicited by licensed inactivated- and live-attenuated influenza vaccines against H3N2 hemagglutinin. Hum Immunol 72:463–9.

33. Tosh PK, Boyce TG, Poland GA. 2008. Flu myths: dispelling the myths associated with live attenuated influenza vaccine. Mayo Clin Proc 83:77–84.

34. Shannon I, White CL, Nayak JL. 2020. Understanding Immunity in Children Vaccinated With Live Attenuated Influenza Vaccine. J Pediatric Infect Dis Soc 9:S10–S14.

35. Sridhar S, Brokstad KA, Cox RJ. 2015. Influenza Vaccination Strategies: Comparing Inactivated and Live Attenuated Influenza Vaccines. Vaccines (Basel) 3:373–89.

36. Darvishian M, Bijlsma MJ, Hak E, van den Heuvel ER. 2014. Effectiveness of seasonal influenza vaccine in community-dwelling elderly people: a meta-analysis of test-negative design case-control studies. Lancet Infect Dis 14:1228–39.

37. Manzoli L, Ioannidis JP, Flacco ME, De Vito C, Villari P. 2012. Effectiveness and harms of seasonal and pandemic influenza vaccines in children, adults and elderly: a critical review and re-analysis of 15 meta-analyses. Hum Vaccin Immunother 8:851–62.

38. Martinez-Sobrido L, Peersen O, Nogales A. 2018. Temperature Sensitive Mutations in Influenza A Viral Ribonucleoprotein Complex Responsible for the Attenuation of the Live Attenuated Influenza Vaccine. Viruses 10.

39. Mohn KG, Brokstad KA, Pathirana RD, Bredholt G, Jul-Larsen A, Trieu MC, Lartey SL, Montemoli E, Tondel C, Aarstad HJ, Cox RJ. 2016. Live Attenuated Influenza Vaccine in Children Induces B-Cell Responses in Tonsils. J Infect Dis 214:722–31.

40. Poehling KA, Caspard H, Peters TR, Belongia EA, Congeni B, Gaglani M, Griffin MR, Irving SA, Kavathekar PK, McLean HQ, Naleway AL, Ryan K, Talbot HK, Ambrose CS. 2018. 2015-2016 Vaccine Effectiveness of Live Attenuated and Inactivated Influenza Vaccines in Children in the United States. Clin Infect Dis 66:665–672.

41. Jang H, Elaish M, Kc M, Abundo MC, Ghorbani A, Ngunjiri JM, Lee CW. 2018. Efficacy and synergy of live-attenuated and inactivated influenza vaccines in young chickens. PLoS One 13:e0195285.

42. Rodriguez L, Nogales A, Reilly EC, Topham DJ, Murcia PR, Parrish CR, Martinez Sobrido L. 2017. A live-attenuated influenza vaccine for H3N2 canine influenza virus. Virology 504:96–106.

43. Nogales A, Rodriguez L, Chauche C, Huang K, Reilly EC, Topham DJ, Murcia PR, Parrish CR, Martinez-Sobrido L. 2017. Temperature-Sensitive Live-Attenuated Canine Influenza Virus H3N8 Vaccine. J Virol 91.

44. Krammer F. 2019. The human antibody response to influenza A virus infection and vaccination. Nat Rev Immunol 19:383–397.

45. Hilimire TA, Nogales A, Chiem K, Ortego J, Martinez-Sobrido L. 2020. Increasing the Safety Profile of the Master Donor Live Attenuated Influenza Vaccine. Pathogens 9.

46. Blanco-Lobo P, Rodriguez L, Reedy S, Oladunni FS, Nogales A, Murcia PR, Chambers TM, Martinez-Sobrido L. 2019. A Bivalent Live-Attenuated Vaccine for the Prevention of Equine Influenza Virus. Viruses 11.

47. Rodriguez L, Reedy S, Nogales A, Murcia PR, Chambers TM, Martinez-Sobrido L. 2018. Development of a novel equine influenza virus live-attenuated vaccine. Virology 516:76–85.

48. Chan W, Zhou H, Kemble G, Jin H. 2008. The cold adapted and temperature sensitive influenza A/Ann Arbor/6/60 virus, the master donor virus for live attenuated influenza vaccines, has multiple defects in replication at the restrictive temperature. Virology 380:304–11.

49. Snyder MH, Betts RF, DeBorde D, Tierney EL, Clements ML, Herrington D, Sears SD, Dolin R, Maassab HF, Murphy BR. 1988. Four viral genes independently contribute to attenuation of live influenza A/Ann Arbor/6/60 (H2N2) cold-adapted reassortant virus vaccines. J Virol 62:488–95.

50. Cox NJ, Kitame F, Kendal AP, Maassab HF, Naeve C. 1988. Identification of sequence changes in the cold-adapted, live attenuated influenza vaccine strain, A/Ann Arbor/6/60 (H2N2). Virology 167:554–67.

51. Cox A, Baker SF, Nogales A, Martinez-Sobrido L, Dewhurst S. 2015. Development of a mouse-adapted live attenuated influenza virus that permits in vivo analysis of enhancements to the safety of live attenuated influenza virus vaccine. J Virol 89:3421–6.

52. Rodriguez L, Nogales A, Murcia PR, Parrish CR, Martinez-Sobrido L. 2017. A bivalent live-attenuated influenza vaccine for the control and prevention of H3N8 and H3N2 canine influenza viruses. Vaccine 35:4374–4381.

53. Nogales A, Rodriguez L, DeDiego ML, Topham DJ, Martinez-Sobrido L. 2017. Interplay of PA-X and NS1 Proteins in Replication and Pathogenesis of a Temperature-Sensitive 2009 Pandemic H1N1 Influenza A Virus. J Virol 91.

54. Cox A, Schmierer J, D’Angelo J, Smith A, Levenson D, Treanor J, Kim B, Dewhurst S. 2020. A Mutated PB1 Residue 319 Synergizes with the PB2 N265S Mutation of the Live Attenuated Influenza Vaccine to Convey Temperature Sensitivity. Viruses 12.

55. Cox A, Dewhurst S. 2015. A Single Mutation at PB1 Residue 319 Dramatically Increases the Safety of PR8 Live Attenuated Influenza Vaccine in a Murine Model without Compromising Vaccine Efficacy. J Virol 90:2702–5.

56. Lowen AC, Bouvier NM, Steel J. 2014. Transmission in the guinea pig model. Curr Top Microbiol Immunol 385:157–83.

57. Lowen AC, Mubareka S, Steel J, Palese P. 2007. Influenza virus transmission is dependent on relative humidity and temperature. PLoS Pathog 3:1470–6.

58. Steel J, Staeheli P, Mubareka S, Garcia-Sastre A, Palese P, Lowen AC. 2010. Transmission of pandemic H1N1 influenza virus and impact of prior exposure to seasonal strains or interferon treatment. J Virol 84:21–6.

59. Song H, Nieto GR, Perez DR. 2007. A new generation of modified live-attenuated avian influenza viruses using a two-strategy combination as potential vaccine candidates. J Virol 81:9238–48.

60. Fan H, Walker AP, Carrique L, Keown JR, Serna Martin I, Karia D, Sharps J, Hengrung N, Pardon E, Steyaert J, Grimes JM, Fodor E. 2019. Structures of influenza A virus RNA polymerase offer insight into viral genome replication. Nature 573:287–290.

61. Noda T, Sugita Y, Aoyama K, Hirase A, Kawakami E, Miyazawa A, Sagara H, Kawaoka Y. 2012. Three-dimensional analysis of ribonucleoprotein complexes in influenza A virus. Nat Commun 3:639.

62. Breen M, Nogales A, Baker SF, Perez DR, Martinez-Sobrido L. 2016. Replication-Competent Influenza A and B Viruses Expressing a Fluorescent Dynamic Timer Protein for In Vitro and In Vivo Studies. PLoS One 11:e0147723.

63. Clark AM, Nogales A, Martinez-Sobrido L, Topham DJ, DeDiego ML. 2017. Functional Evolution of Influenza Virus NS1 Protein in Currently Circulating Human 2009 Pandemic H1N1 Viruses. J Virol 91.

64. DiPiazza A, Nogales A, Poulton N, Wilson PC, Martinez-Sobrido L, Sant AJ. 2017. Pandemic 2009 H1N1 Influenza Venus reporter virus reveals broad diversity of MHC class II-positive antigen-bearing cells following infection in vivo. Sci Rep 7:10857.

65. Nogales A, Martinez-Sobrido L, Chiem K, Topham DJ, DeDiego ML. 2018. Functional Evolution of the 2009 Pandemic H1N1 Influenza Virus NS1 and PA in Humans. J Virol 92.

66. Guo H, Santiago F, Lambert K, Takimoto T, Topham DJ. 2011. T cell-mediated protection against lethal 2009 pandemic H1N1 influenza virus infection in a mouse model. J Virol 85:448–55.

67. Nogales A, Baker SF, Ortiz-Riano E, Dewhurst S, Topham DJ, Martinez-Sobrido L. 2014. Influenza A virus attenuation by codon deoptimization of the NS gene for vaccine development. J Virol 88:10525–40.

68. Nogales A, Baker SF, Martinez-Sobrido L. 2014. Replication-competent influenza A viruses expressing a red fluorescent protein. Virology 476C:206–216.

69. Baker SF, Nogales A, Finch C, Tuffy KM, Domm W, Perez DR, Topham DJ, Martinez-Sobrido L. 2014. Influenza A and B virus intertypic reassortment through compatible viral packaging signals. J Virol 88:10778–91.

70. Chiem K, Lopez-Garcia D, Ortego J, Martinez-Sobrido L, DeDiego ML, Nogales A. 2021. Identification of amino acid residues required for inhibition of host gene expression by influenza A/Viet Nam/1203/2004 H5N1 PA-X. J Virol:JVI0040821.

71. Cheng BY, Ortiz-Riano E, Nogales A, de la Torre JC, Martinez-Sobrido L. 2015. Development of live-attenuated arenavirus vaccines based on codon deoptimization. J Virol 89:3523–33.

72. Nakajima Y, Kobayashi K, Yamagishi K, Enomoto T, Ohmiya Y. 2004. cDNA cloning and characterization of a secreted luciferase from the luminous Japanese ostracod, Cypridina noctiluca. Biosci Biotechnol Biochem 68:565–70.

73. Liu WC, Nachbagauer R, Stadlbauer D, Solorzano A, Berlanda-Scorza F, Garcia-Sastre A, Palese P, Krammer F, Albrecht RA. 2019. Sequential Immunization With Live-Attenuated Chimeric Hemagglutinin-Based Vaccines Confers Heterosubtypic Immunity Against Influenza A Viruses in a Preclinical Ferret Model. Front Immunol 10:756.

